# Population dynamics of multi-host communities attacked by a common parasitoid

**DOI:** 10.1101/2021.01.04.425210

**Authors:** Abhyudai Singh

## Abstract

We model population dynamics of two host species attacked by a common parasitoid using a discrete-time formalism that captures their population densities from year to year. It is well known starting from the seminal work of Nicholson and Bailey that a constant parasitoid attack rate leads to an unstable host-parasitoid interaction. However, a Type III functional response, where the parasitoid attack rate accelerates with increasing host density stabilizes the population dynamics. We first consider a scenario where both host species are attacked by a parasitoid with the same Type III functional response. Our results show that sufficient fast acceleration of the parasitoid attack rate stabilizes the population dynamics of all three species. For two symmetric host species, the extent of acceleration needed to stabilize the three-species equilibrium is exactly the same as that needed for a single host-parasitoid interaction. However, asymmetry can lead to scenarios where the removal of a host species from a stable interaction destabilizes the interaction between the remaining host species and the parasitoid. Next, we consider a situation where one of the host species is attacked at a constant rate (i.e., Type I functional response), and the other species is attacked via a Type III functional response. We identify parameter regimes where a Type III functional response to just one of the host species stabilizes the three species interaction. In summary, our results show that a generalist parasitoid with a Type III functional response to one or many host species can play a key role in stabilizing population dynamics of host-parasitoid communities in apparent competition.

## I. Introduction

The population dynamics of host-parasitoid interactions has been traditionally modeled using a discrete-time formalism starting from the classical Nicholson-Bailey model

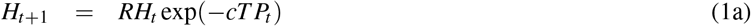

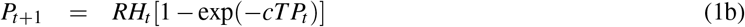

where *H*_*t*_ and *P*_*t*_ are the adult host, and the adult parasitoid densities, respectively, at the start of year *t* [1]–[8]. The model is motivated by the annual insect life cycles living in the temperate regions of the world that consists of adult hosts emerging during spring, laying eggs that hatch into larvae. Host larvae then overwinter in the pupal stage, and metamorphosize as adults the following year. The host becomes vulnerable to parasitoid attacks in one of its developmental stages, and for the sake of convenience, we assume this to be the host larval stage. Adult female parasitoids emerge during spring, search and attack hosts by laying an egg into the body of the host. While adult parasitoids die after this time window, the parasitoid egg hatches into a juvenile parasitoid that grows at the host’s expense by using it as a food source, ultimately resulting in the death of the host [9]–[12]. The juvenile parasitoids pupate, overwinter, and emerge as adult parasitoids the following year. In (1), *RH*_*t*_ is the host larval density exposed to parasitoid attacks at the start of the vulnerable stage, where *R* > 1 denotes the number of viable eggs produced by each adult host. Assuming parasitoids attack and parasitize hosts at a constant rate *c*, results in exp(−*cT P*_*t*_) fraction of host larvae escaping parasitism, where *T* is the duration of the host vulnerable stage [2], [3], [13]. Thus, *RH*_*t*_ exp(−*cT P*_*t*_) is the total larval density escaping parasitism to become adult hosts for next year. Finally, *RH*_*t*_[1 − exp(−*cT P*_*t*_)] is the density of parasitized larvae that give rise to adult parasitoids in the next generation.

It turns out that the Nicholson-Bailey model is characterized by diverging oscillations in population densities resulting in an unstable population dynamics [2]. A simple generalization of (1) is

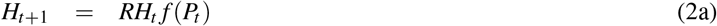

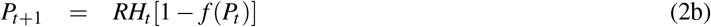

where the fraction host larvae escaping parasitism is represented by a monotonically decreasing function *f* that only depends on the parasitoid density [3], [14], [15]. As has been done in the literature, we refer to *f* as the *escape response*. For (2), the unique non-trivial host-parasitoid equilibrium *H*\* and *P*\* is given as the solution to

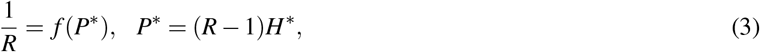

and this equilibrium is stable, if and only, if,

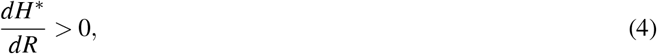

i.e., the function *f* leads to an equilibrium adult host density *H*\* that is an *increasing* function of the host reproduction rate *R* [16], [17]. Many types of population interactions and ecological factors lead to models of the form (2) and stabilize host-parasitoid interactions. Some examples include, a fraction of the host population being in a refuge (i.e., protected from parasitoid attacks) [3], [18], large host-to-host difference in parasitism risk [16], [19]–[21], parasitoid interference [22]–[24], and aggregation in parasitoid attacks [25]–[27].

Model (2) can be expanded to capture the population dynamics of two different host species H and Q attacked by the same parasitoid

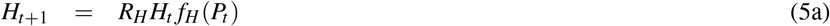

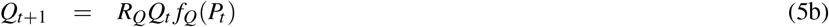

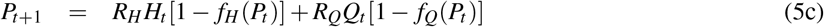

where *H*_*t*_ and *Q*_*t*_ represent the densities of the two host species with reproduction rate *R*_*H*_, *R*_*Q*_, and parasitoid-dependent escape responses *f*_*H*_, *f*_*Q*_, respectively. This indirect interaction between two hosts (due to a shared parasitoid) has been referred to as “apparent competition” [28]–[32] and well documented in field observations [33]–[35]. Even if each host species by itself has a stable equilibrium with the parasitoid, stable coexistence of *both* host species is not possible in (5) with apparent competition driving the host species with the lower reproduction rate to extinction [31]. This leads to an intriguing question: can a host-density dependent escape response in (5) yield stable coexistence of all three species? To this end, we systematically explore the role of a Type III parasitoid functional response in regulating the population dynamics of apparent competition. While much literature on host-parasitoid interactions points to Type I and II functional responses [36]–[40], there has been some empirical evidence for a Type III response [41]–[43]. Functional response can have important implications for biological control of pests by parasitoids [44]–[48], and a Type III functional response has been shown to suppress the host density to arbitrary low levels while maintaining system stability [48]. We start by reviewing recent work that incorporates a Type III functional response in discrete-time models using a semi-discrete or hybrid framework.

## II. Type III functional response results in a host-density dependent escape response

In our prior work, we have considered a Type III parasitoid functional response, where the attack rate *cL*^*m*^ accelerates with increasing host larvae density *L* for some positive constant *c* and exponent *m*. Here, *L* denotes the non-parasitized larval density that decreases over time during the vulnerable stage leading to a variable attack rate. To capture such effects of populations changing continuously within the larval stage of each year, a semi-discrete or hybrid formalism has been proposed to mechanistically formulate the corresponding discrete-time model [13], [48]–[52]. We briefly describe this semi-discrete approach and illustrate its application in formulating a Type III functional response.

Let *τ* denote the time within the host vulnerable stage that varies from 0 to *T* corresponding to the start and end of the vulnerable stage. The densities of parasitoids, un-parasitized and parasitized host larvae at time *τ* within the vulnerable stage of year *t* are represented by *P*(*τ, t*), *L*(*τ, t*), *I*(*τ, t*), respectively. These densities evolve as per the dynamical system

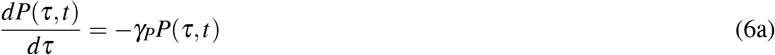

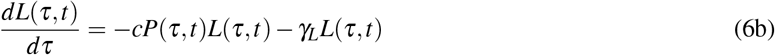

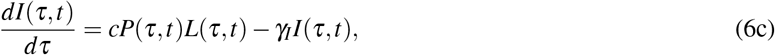

where *c* represents the parasitoid’s attack rate *per host*, and *γ*_*P*_, *γ*_*L*_, *γ*_*I*_ are the death rates of the respective species. Assuming *P*_*t*_ parasitoids, *RH*_*t*_ host larvae, and no parasitized larvae at the start of the vulnerable period (*τ* = 0), solving the above differential equations with initial conditions

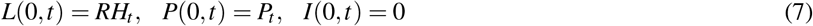

predicts the parasitized and unparasitized larval populations at the end of the season (*τ* = *T*). This leads to a more general discrete-time model

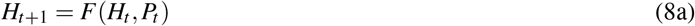

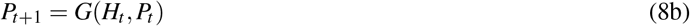

where update functions are obtained by setting

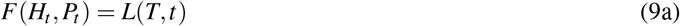

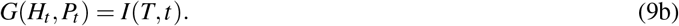

Solving (6) for a constant attack rate *c* with no mortalities (*γ*_*P*_ = *γ*_*L*_ = *γ*_*I*_ = 0)

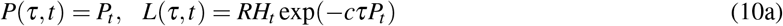

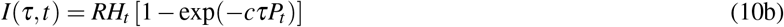

which using (9) yields the Nicholson-Bailey model (1).

To model a Type III parasitoid functional response, we substitute the attack rate *c* in (6) with *cL*(*τ, t*)^*m*^ which leads to the model (assuming without loss of generality *T* = 1)

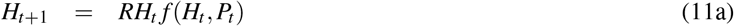

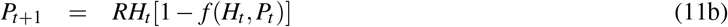

with an escape response

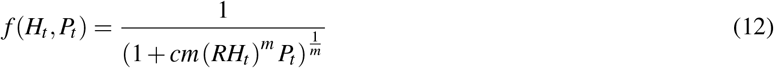

that depends on both host and parasitoid population densities [13]. It turns out that the model’s unique non-trivial fixed point

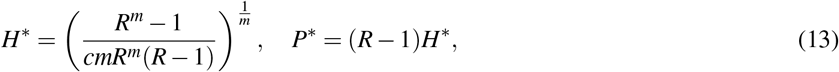

is stable iff *m* > 1, and *m* = 1 results in a neutrally stable equilibrium where populations oscillate with a period of 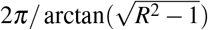 [13]. Interestingly, in contrast to the stability condition (4) that arise for parasitoid-dependent escape responses, here *H*\* is a decreasing function of *R*. This contrasting behaviors of *H*\* with respect to *R* can be exploited for discriminating between stability mechanisms [53]. It is important to point out that a phenomenological approach of incorporating a Type III functional response by simply substituting *c* in the Nicholson-bailey model (1) with *c*(*RH*_*t*_)^*m*^ (i.e., the parasitoid attack rate is set by the initial larval density *RH*_*t*_ and remains fixed through the larval stage) leads to an unstable population equilibrium for all *m* ≥ 0 [54], [55]. Finally, we point out that here we having ignored saturation effects in the attack rate with increasing host larval density, and refer the reader to [13] for a more general treatment of Type III functional response.

## III. Population dynamics of apparent competition

Having introduced a discrete-time formulation of a Type III functional response, we now consider two scenarios of apparent competition: i) both host species are attacked by a common parasitoid with the same Type III functional response; ii) the parasitoid attacks one host species at a constant rate (as in the Nicholson-Bailey model) but exhibits a Type III functional response for the other host. We consider the former case first.

### A. Type III functional response to both host species

Let the parasitoid attack the two host species with rates 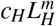 and 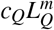, where *L*_*H*_ and *L*_*Q*_ denote the larval densities of hosts H and Q, respectively. Here, *m* quantifies the extent of Type III functional response and is assumed to be the same for both hosts. Then, the apparent competition can be modeled as

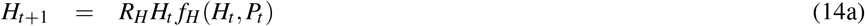

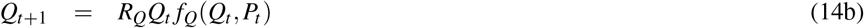

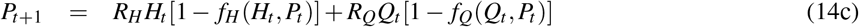

with escape responses

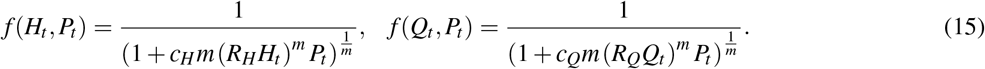

This model has a unique non-trivial equilibrium given by

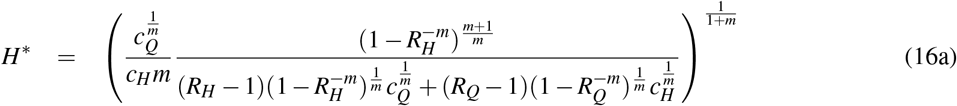

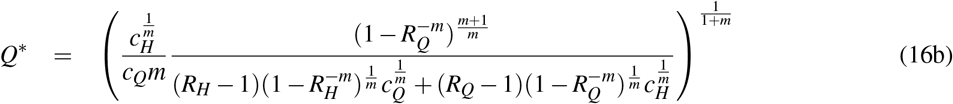

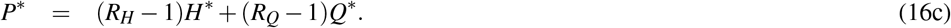

To perform a local stability analysis, we consider small perturbations *H*_*t*_, *q*_*t*_ and *p*_*t*_

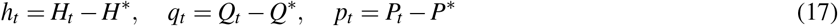

around the fixed point (16) that results in the following *linear* discrete-time system

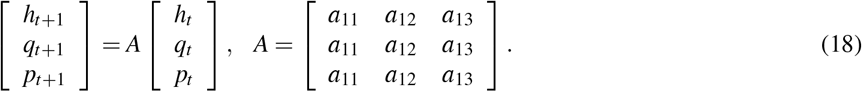

The entries of the Jacobian matrix *A* are given by

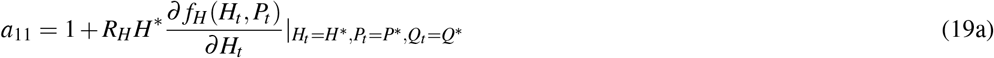

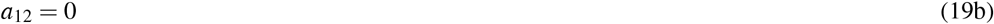

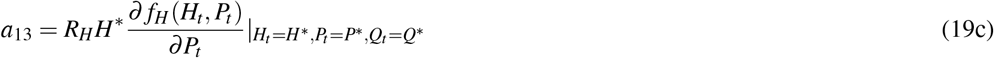

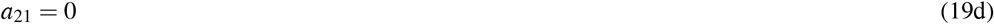

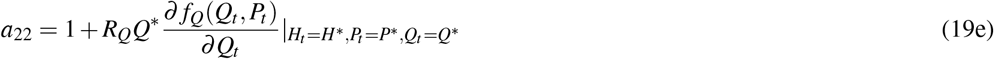

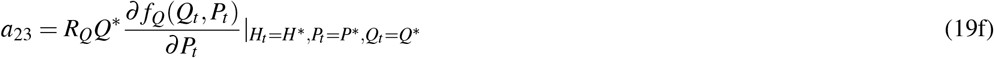

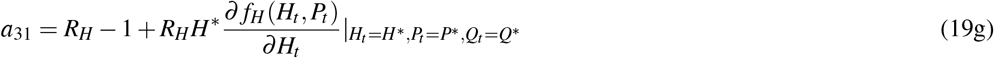

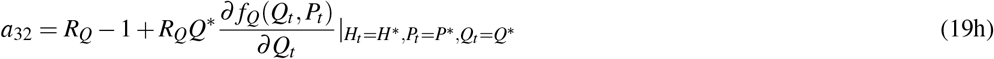

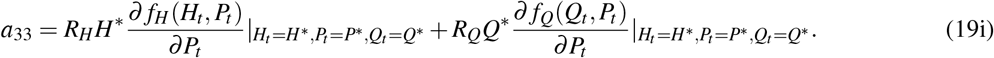

where

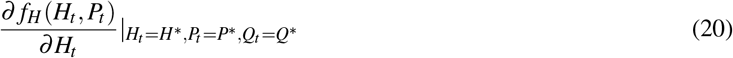

denotes the partial derivative of *f*_*H*_ with respect to *H*_*t*_ evaluated at the equilibrium point.

The stability of the equilibrium point requires all eigenvalues of *A* to be inside the unit circle, i.e., have an absolute value less than one [56], [57]. For a 3 × 3 matrix, this corresponds to the following result. Let

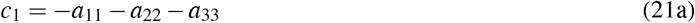

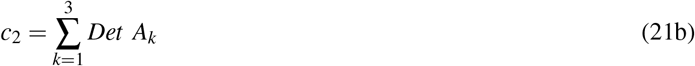

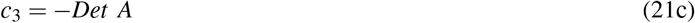

where *A*_*k*_ is the 2 × 2 matrix obtained from matrix *A* by deleting row *k* and column *k*, and *Det* represents the matrix determinant. Then, the non-trivial fixed point *H*\*, *Q*\*, *P*\* is asymptotically stable, if and only if, the following inequalities hold

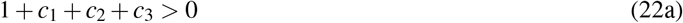

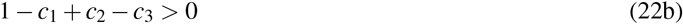

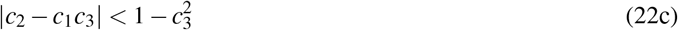

[56], [57]. Our analysis of inequalities in Mathematica [58] reveal that here stability is determined by (22c), and for symmetric host reproduction rates *R*_*H*_ = *R*_*Q*_ the equilibrium (16) is stable, iff, *m* > 1. Recall from the previous section that this condition is identical to that required for stability of a single host-parasitoid interaction. Intriguingly, asymmetric hosts where *R*_*H*_ ≠ *R*_*Q*_ and *c*_*H*_ ≠ *c*_*Q*_ can lead to scenarios where stability of the three-species interaction can occur even when *m* < 1. This is illustrated in Fig. 1 which plots the critical value of *m* needed for stability as a function of *R*_*Q*_/*R*_*H*_ and *c*_*Q*_/*c*_*H*_. For example, when *R*_*Q*_/*R*_*H*_ = 8 and *c*_*Q*_/*c*_*H*_ = 30, the system is stable for *m* > 0.8. As expected, when stable coexistence of both hosts occurs for values of *m* < 1, then removal of one of the hosts results in an unstable population dynamics (Fig. 2).

**Fig. 1:**
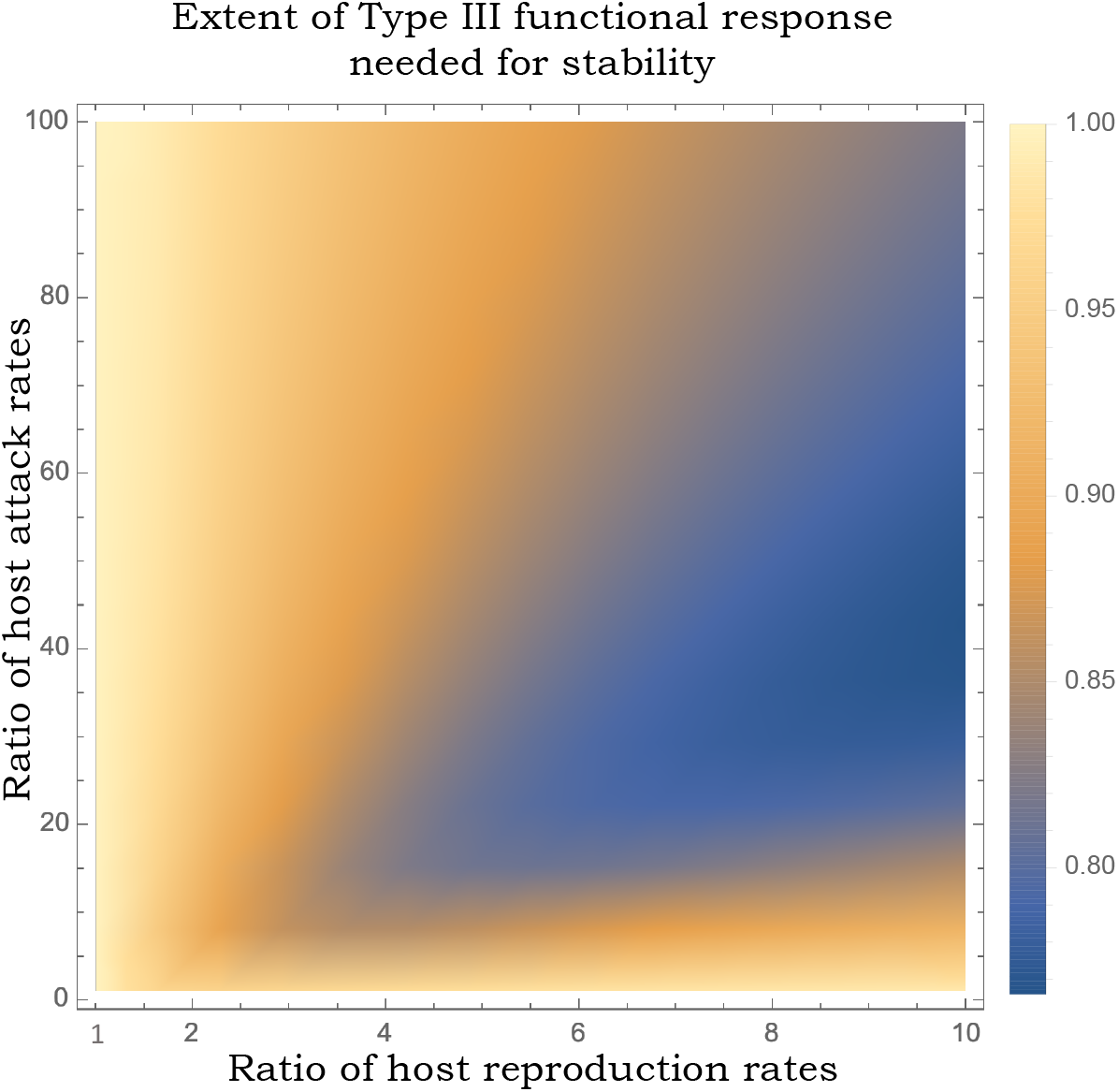
The critical value of *m* needed to stabilize the equilibrium (16) in the apparent competition model (14) as a function of *R*_*Q*_/*R*_*H*_ and *c*_*Q*_/*c*_*H*_. When *R*_*H*_ = *R*_*Q*_ the equilibrium (16) is stable, iff, *m* > 1, while for large values of *R*_*Q*_/*R*_*H*_ and *c*_*Q*_/*c*_*H*_ system stability can arise for *m* < 1. In this plot, we assume *R*_*H*_ = 2 and *c*_*H*_ = 1.

**Fig. 2:**
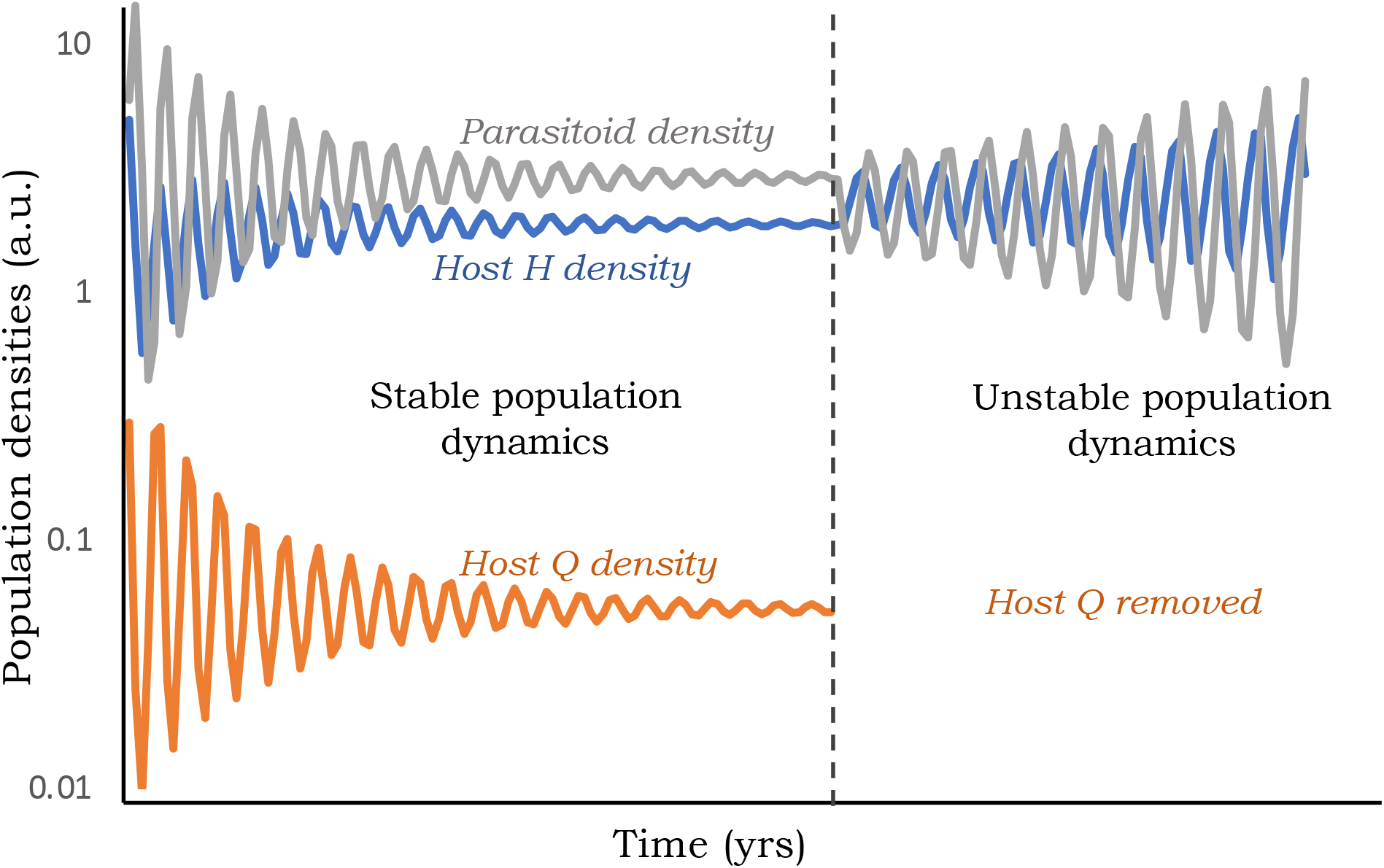
Simulation of model (14) for *R*_*H*_ = 2, *R*_*Q*_ = 16, *c*_*H*_ = 1, *c*_*Q*_ = 30 and *m* = 0.9. This leads to a stable population dynamics with coexistence of both host species in apparent competition due to the shared natural enemy. For *m* = 0.9, removal of host Q results in an unstable interaction with diverging oscillations as stability for a single host-parasitoid interaction requires *m* > 1.

### B. Type III functional response to just one host species

We next consider the model

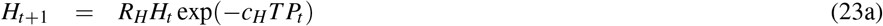

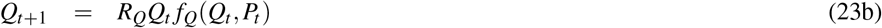

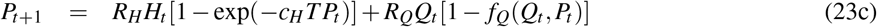

which corresponds to the parasitoid attacking host H with a constant rate *c*_*H*_ as in the Nicholson-Bailey model (i.e., a Type I functional response), and attacking host Q with a Type III functional response which leads to

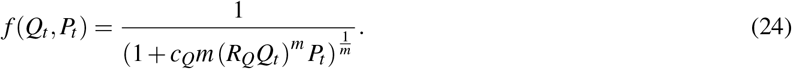

Model (23) can be further expanded by relaxing a key assumption in the Nicholson-Bailey model formulation. In particular, the Nicholson-Bailey model assumes that all hosts are identical in terms of their vulnerability to parasitism, and host-to-host differences in parasitism risk have been modeled in literature by replacing *c*_*H*_*T* in (23a) by a Gamma-distributed random variable with mean 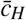 and coefficient of variation *CV* [16], [20], [59]–[62]. Taking the expected value of the escape response exp(−*c*_*H*_*T P*_*t*_) transforms model (23) into

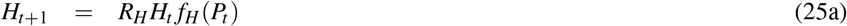

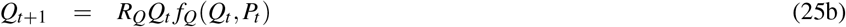

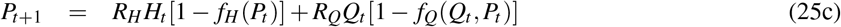

where

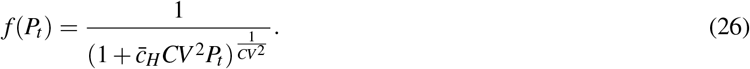

This leads to a scenario where host H has a parasitoid-dependent escape response arising from variations in parasitism risk, while host Q’s escape response depends on both host and parasitoid densities due to the Type III functional response. Each host species *by itself* will have a stable equilibrium with the parasitoid if *CV* > 1 for H [19], [21], [27] (in the absence of Q), and *m* > 1 for Q (in the absence of H). *For what values of CV and m can both H and Q coexist?*

The non-trivial fixed point of (25) is given by

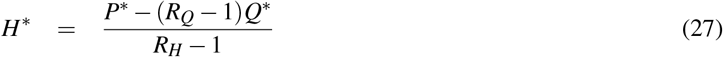

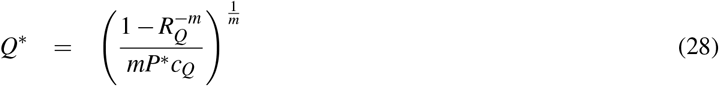

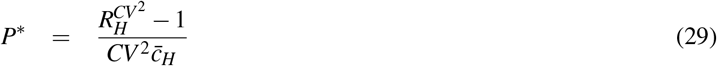

and *H*\* > 0 requires the parasitoid attack rate to H be sufficient small

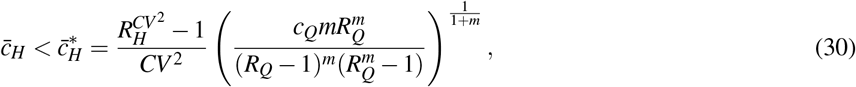

without which host *H* will be driven to extinction. Linearization of the model (25) around the above equilibrium results in the Jacobian matrix *A* (19) where now ∂ *f*_*H*_(*H*_*t*_, *P*_*t*_)/∂*H*_*t*_ = 0. The stability region as predicted by inequalities (22) is graphically illustrated in Fig. 3 and reveals:

1) When both *CV, m* > 1 (i.e., each host species by itself has a stable interaction with the parasitoid), stable coexistence of both host species and the parasitoid is always possible.
2) Significant regions of coexistence of both host species and the parasitoid involve either one of *CV* or *m* being larger than one. Here, removal of the corresponding host (for example, removal of host *Q* when *m* > 1) will destabilize the single-host parasitoid interaction.
3) Interestingly, there is a small region where coexistence occurs when both *CV, m* < 1. Here, each host by itself has an unstable equilibrium with the parasitoid, and stability of the three-species equilibrium occurs with apparent competition.
4) When *CV* = 0, i.e. host H is attacked at a constant rate, then sufficient strong acceleration in the Type III response towards host Q leads to stable coexistence of both H and Q. This critical value of *m* ≈ 1.15 when *R*_*H*_ = *R*_*Q*_ = 2 and *m* ≈ 1.6 when *R*_*H*_ = *R*_*Q*_ = 10.

**Fig. 3:**
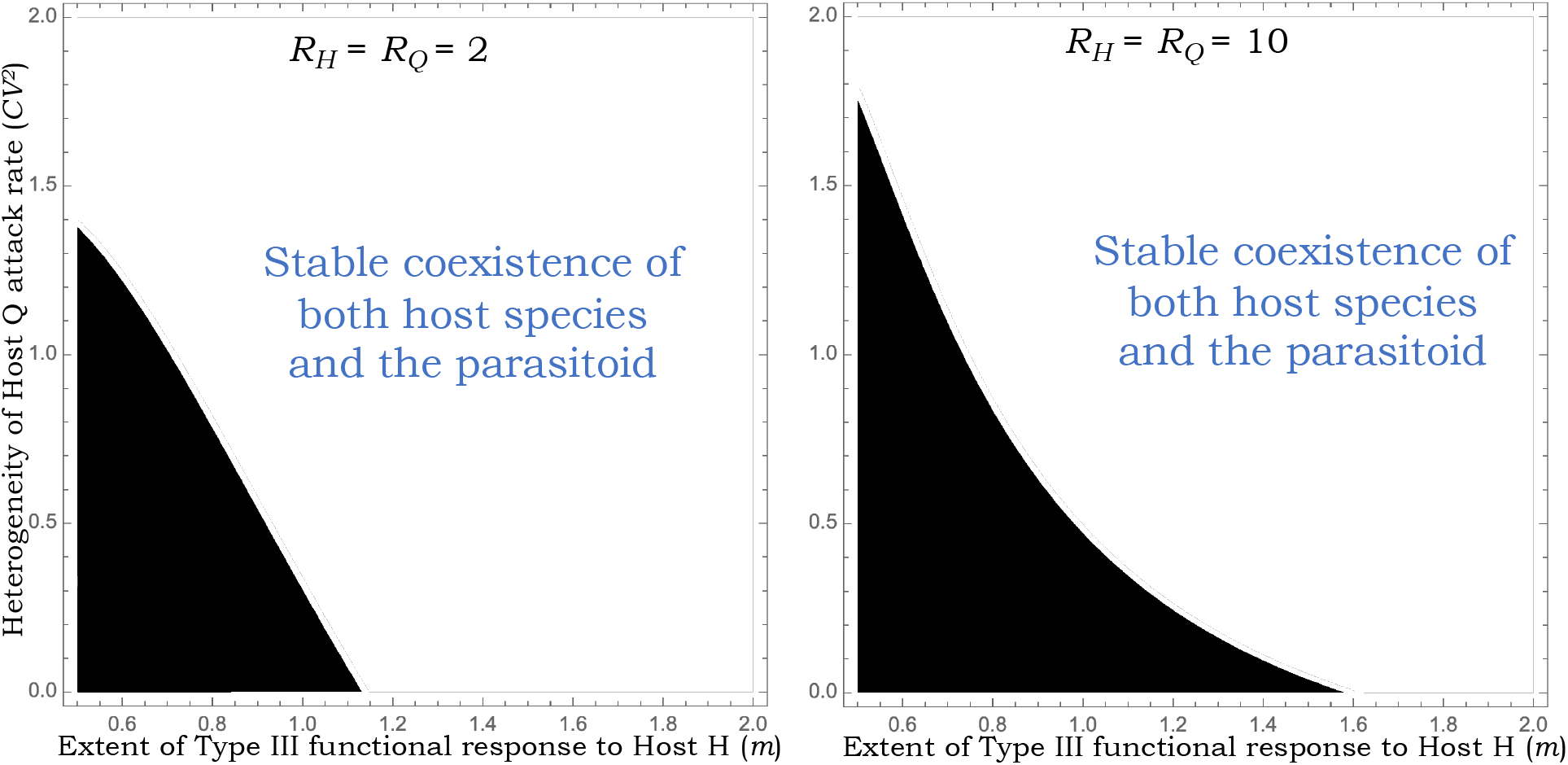
Stability regions for the equilibrium point (27) of the apparent competition model (25) as a function of *CV* and *m* for *R*_*H*_ = *R*_*Q*_ = 2 (left) and *R*_*H*_ = *R*_*Q*_ = 10 (right). The black shaded (unshaded) regions denote the equilibrium being unstable (stable). From (30), existence of the equilibrium point (27) requires 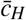 to be below a critical value 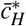 and for this plot we assume 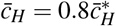.

In summary, these results strongly argue that generalist parasitoids attacking multiple host species play an important role in stabilizing the overall population dynamics. The coexistence of host species in apparent competition due to a shared natural enemy requires parasitoids to have sigmoidal functional responses towards one or multiple species with attack rates accelerating in response to host density.

## Notes

### Competing Interest Statement

The authors have declared no competing interest.

## REFERENCES

1. A. Singh, “Analytical discrete-time models of insect population dynamics,” 2020.

2. A. Nicholson and V. A. Bailey, “The balance of animal populations. part 1.” Proc. of Zoological Society of London, vol. 3, pp. 551–598, 1935.

3. W. W. Murdoch, C. J. Briggs, and R. M. Nisbet, Consumer-Resouse Dynamics. Princeton, NJ: Princeton University Press, 2003.

4. L. Edelstein-Keshet, Mathematical models in biology. SIAM, 2005.

5. N. Kakehashi, Y. Suzuki, and Y. Iwasa, “Niche overlap of parasitoids in host-parasitoid systems: its consequence to single versus multiple introduction controversy in biological control,” Journal of Applied Ecology, pp. 115–131, 1984.

6. R. M. May and M. P. Hassell, “The dynamics of multiparasitoid-host interactions,” The American Naturalist, vol. 117, no. 3, pp. 234–261, 1981.

7. E. Hackett-Jones, C. Cobbold, and A. White, “Coexistence of multiple parasitoids on a single host due to differences in parasitoid phenology,” Theoretical Ecology, vol. 2, no. 1, pp. 19–31, 2009.

8. E. van Velzen, S. Pérez-Vila, and R. S. Etienne, “The role of within-host competition for coexistence in multiparasitoid-host systems,” The American Naturalist, vol. 187, no. 1, pp. 48–59, 2016.

9. A. E. Hajek, Insect parasitoids: attack by aliens. Cambridge University Press, 2004, p. 145169.

10. H. C. J. Godfray, Parasitoids; Behavioral and Evolutionary Ecology. 41 William St, Princeton, NJ 08540: Princeton University Press, 1994.

11. J. Waage and D. Greathead, Insect Parasitoids. Academic Press, 1986.

12. M. E. Hochberg and A. R. Ives, Parasitoid population biology. Princeton University Press, 2000.

13. A. Singh and R. M. Nisbet, “Semi-discrete host-parasitoid models,” Journal of Theoretical Biology, vol. 247, no. 4, pp. 733–742, 2007.

14. M. P. Hassell. New York: Oxford University Press, 2000.

15. W. S. C. Gurney and R. M. Nisbet, Ecological Dynamics. Oxford University Press, 1998.

16. A. Singh, W. W. Murdoch, and R. M. Nisbet, “Skewed attacks, stability, and host suppression,” Ecology, vol. 90, no. 6, pp. 1679–1686, 2009.

17. A. Singh, “Generalized conditions for coexistence of competing parasitoids on a shared host,” bioRxiv, 2020.

18. E. Bešo, S. Kalabušić, N. Mujić, and E. Pilav, “Stability of a certain class of a host–parasitoid models with a spatial refuge effect,” Journal of Biological Dynamics, vol. 14, no. 1, pp. 1–31, 2020.

19. A. D. Taylor, “Heterogeneity in host-parasitoid interactions: ’aggregation of risk’ and the ’*cv*^2^ > 1 rule.’,” Trends in Ecology and Evolution, vol. 8, pp. 400–405, 1993.

20. M. P. Hassell, R. M. May, S. W. Pacala, and P. L. Chesson., “The persistence of host–parasitoid associations in patchy environments. I. a general criterion.” American Naturalist, vol. 138, pp. 568–583, 1991.

21. S. W. Pacala and M. P. Hassell., “The persistence of host– parasitoid associations in patchy environments. II. evaluation of field data.” American Naturalist, vol. 138, pp. 584–605, 1991.

22. C. Bernstein, “Density dependence and the stability of host-parasitoid systems,” Oikos, pp. 176–180, 1986.

23. C. Free, J. Beddington, and J. Lawton, “On the inadequacy of simple models of mutual interference for parasitism and predation,” The Journal of Animal Ecology, pp. 543–554, 1977.

24. D. Rogers and M. Hassell, “General models for insect parasite and predator searching behaviour: interference,” The Journal of Animal Ecology, pp. 239–253, 1974.

25. J. D. Reeve, J. T. Cronin, and D. R. Strong., “Parasitoid aggregation and the stabilization of a salt marsh host– parasitoid system,” Ecology, vol. 75, pp. 288–295, 1994.

26. P. Rohani, H. C. J. Godfray, and M. P. Hassell, “Aggregation and the dynamics of host-parasitoid systems: A discrete-generation model with within-generation redistribution,” The American Naturalist, vol. 144, no. 3, pp. 491–509, 1994.

27. R. M. May, “Host–parasitoid systems in patchy environments: a phenomenological model,” Journal of Animal Ecology, vol. 47, pp. 833–844, 1978.

28. R. D. Holt and M. B. Bonsall, “Apparent competition,” Annual Review of Ecology, Evolution, and Systematics, vol. 48, pp. 447–471, 2017.

29. S. J. Schreiber, “The 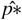 rule in the stochastic holt-lawton model of apparent competition,” arXiv preprint arXiv:2007.01953, 2020.

30. P. Hudson and J. Greenman, “Competition mediated by parasites: biological and theoretical progress,” Trends in Ecology & Evolution, vol. 13, no. 10, pp. 387–390, 1998.

31. R. D. Holt and J. H. Lawton, “Apparent competition and enemy-free space in insect host-parasitoid communities,” The American Naturalist, vol. 142, no. 4, pp. 623–645, 1993.

32. M. B. Bonsall and M. P. Hassell, “Parasitoid-mediated effects: apparent competition and the persistence of host–parasitoid assemblages,” Researches on Population Ecology, vol. 41, no. 1, pp. 59–68, 1999.

33. C. M. Frost, G. Peralta, T. A. Rand, R. K. Didham, A. Varsani, and J. M. Tylianakis, “Apparent competition drives community-wide parasitism rates and changes in host abundance across ecosystem boundaries,” Nature communications, vol. 7, no. 1, pp. 1–12, 2016.

34. C. a. Muller and H. Godfray, “Apparent competition between two aphid species,” Journal of Animal Ecology, pp. 57–64, 1997.

35. T. Hirao and M. Murakami, “Quantitative food webs of lepidopteran leafminers and their parasitoids in a japanese deciduous forest,” Ecological Research, vol. 23, no. 1, pp. 159–168, 2008.

36. C. S. Holling, “The functional response of predators to prey density and its role in mimicry and population regulation,” The Memoirs of the Entomological Society of Canada, vol. 97, no. S45, pp. 5–60, 1965.

37. N. Mills and I. Lacan, “Ratio dependence in the functional response of insect parasitoids: evidence from trichogramma minutum foraging for eggs in small host patches,” Ecological Entomology, vol. 29, no. 2, pp. 208–216, 2004.

38. X. Chen, S. W. Wong, and P. A. Stansly, “Functional response of tamarixia radiata (hymenoptera: Eulophidae) to densities of its host, diaphorina citri (hemiptera: Psylloidea),” Annals of the Entomological Society of America, vol. 109, no. 3, pp. 432–437, 2016.

39. G. Kaçar, X.-G. Wang, A. Biondi, and K. M. Daane, “Linear functional response by two pupal drosophila parasitoids foraging within single or multiple patch environments,” PLoS One, vol. 12, no. 8, p. e0183525, 2017.

40. J. Ebrahimifar, A. Jamshidnia, and H. Allahyari, “Functional response of eretmocerus delhiensis on trialeurodes vaporariorum by parasitism and host feeding,” Journal of Insect Science, vol. 17, no. 2, p. 56, 2017.

41. M. Collins, S. Ward, and A. Dixon, “Handling time and the functional response of aphelinus thomsoni, a predator and parasite of the aphid drepanosiphum platanoidis,” The Journal of Animal Ecology, pp. 479–487, 1981.

42. M. Hassell, J. Lawton, and J. Beddington, “Sigmoid functional responses by invertebrate predators and parasitoids,” The Journal of Animal Ecology, pp. 249–262, 1977.

43. V. Fernández-arhex and J. C. Corley, “The functional response of parasitoids and its implications for biological control,” Biocontrol Science and Technology, vol. 13, no. 4, pp. 403–413, 2003.

44. S. D. Lane, C. M. St. Mary, and W. M. Getz, “Coexistence of attack-limited parasitoids sequentially exploiting the same resource and its implications for biological control,” in Annales Zoologici Fennici. JSTOR, 2006, pp. 17–34.

45. B. S. Pedersen and N. J. Mills, “Single vs. multiple introduction in biological control: the roles of parasitoid efficiency, antagonism and niche overlap,” Journal of Applied Ecology, vol. 41, no. 5, pp. 973–984, 2004.

46. P. K. Abram, J. Brodeur, V. Burte, and G. Boivin, “Parasitoid-induced host egg abortion; an underappreciated component of biological control services provided by egg parasitoids.” Biological Control, no. 98, pp. 52–60, 2016.

47. M. A. Jervis, B. A. Hawkin, and N. A. C. Kidd, “The usefulness of destructive host-feeding parasitoids in classical biological control: theory and observation conflict,” Ecological Entomology, vol. 21, no. 1, pp. 41–46, 1996.

48. A. Singh and B. Emerick, “Hybrid systems modeling of ecological population dynamics,” bioRxiv, 2020.

49. A. Singh and R. M. Nisbet, “Variation in risk in single-species discrete-time models,” Mathematical Biosciences and Engineering, vol. 5, pp. 859–875, 2008.

50. B. K. Emerick and A. Singh, “The effects of host-feeding on stability of discrete-time host-parasitoid population dynamic models.” Mathematical Biosciences, vol. 272, pp. 54–63, 2016.

51. E. Pachepsky, R. M. Nisbet, and W. W. Murdoch, “Between discrete and continuous: Consumer-resource dynamics with synchronized reproduction,” Ecology, vol. 89, no. 1, pp. 280–288, 2007.

52. B. K. Emerick and A. Singh, “Global redistribution and local migration in semi-discrete host-parasitoid population dynamic models.” Mathematical Biosciences, vol. 327, p. 108409, 2020.

53. A. Singh, “Fluctuations in population densities inform stability mechanisms in host-parasitoid interactions,” bioRxiv, 2020.

54. D. J. Rogers, “Random searching and incest population models,” J. of Animal Ecology, vol. 41, pp. 369–383, 1972.

55. M. P. Hassell and H. N. Comins, “Sigmoid functional responses and population stability,” Theoretical Population Biology, vol. 14, pp. 62–66, 1978.

56. G. Ledder, Mathematics for the life sciences: calculus, modeling, probability, and dynamical systems. Springer Science & Business Media, 2013.

57. S. Elaydi, An Introduction to Difference Equations. Newyork: Springer, 1996.

58. W. R. Inc., “Mathematica, Version 12.2.” [Online]. Available: https://www.wolfram.com/mathematica

59. T. Okuyama, “Density-dependent distribution of parasitism risk among underground hosts.” Bulletin of Entomological Research, vol. 109, no. 4, pp. 528–533, 2019.

60. C. A. Cobbold, J. Roland, and M. A. Lewis, “The impact of parasitoid emergence time on host-parastioid population dynamics,” Theor Popul Biol, vol. 75, no. 2, pp. 201–215, 2009.

61. H. Liere, D. Jackson, and J. Vandermeer, “Ecological complexity in a coffee agroecosystem: spatial heterogeneity, popoulation persistence and biological control,” PLoS One, vol. 7, no. 9, 2012.

62. N. Zoroa, E. Lesigne, M. J. Fernandez-Saez, P. Zoroa, and J. Casas, “The coupon collector urn model with unequal probabilities in ecology and evolution,” Journal of The Royal Society Interface, vol. 14, no. 127, 2017.

